# Ploidy reorganizes ionomic composition across metabolically active and mineralized tissues

**DOI:** 10.64898/2026.02.26.708366

**Authors:** Punidan D. Jeyasingh, Marissa Roseman, Josh Bliss, Yetkin Ipek, Maurine Neiman

## Abstract

Genome size, defined as total nuclear DNA content, varies widely within species and can influence cell size, metabolic scaling, and organismal performance. Yet the mechanisms linking genome architecture to phenotype remain unresolved, especially in animals. Because organisms are far-from-equilibrium systems built from interacting chemical elements, genome size variation should reorganize elemental allocation across tissues rather than shift single-element requirements in isolation. We tested this hypothesis in the New Zealand mudsnail (*Potamopyrgus antipodarum*), which includes co-occurring diploid, triploid, and tetraploid individuals and exhibits strong compartmentalization between metabolically active soft tissue and mineralized shell. We quantified multielemental composition of both tissues and analyzed allocation using additive log-ratios anchored to the measured elemental pool. Tissue type explained most multielemental variance, confirming distinct ionomic regimes for shell and soft tissue. Across tissues, ploidy was associated with significant redistribution of relative elemental allocation despite weak single-element effects. Ploidy-associated imbalances were more pronounced in shell than soft tissue, consistent with long-term integration of elemental fluxes in inert structures and buffering in active tissues. Genome size variation therefore reshapes organismal chemistry through coordinated multielemental redistribution, linking genome architecture to system-level chemical organization.

## Introduction

Genome size, defined here as total nuclear genomic DNA content, varies widely within and among species and can influence cell size, gene dosage, and physiological scaling (Cavalier-Smith 2005; Gregory 2005; Otto 2007; Leitch & Leitch 2008; Roddy et al. 2020; Itgen et al. 2022). Polyploidy provides a tractable form of genome size variation because it generates discrete, heritable differences in nuclear DNA content that can reshape organismal performance and life-history traits (Otto 2007). Despite extensive study, however, the mechanisms linking genome architecture to organismal phenotype remain incompletely resolved, particularly in animals, where polyploidy is less common and often context dependent (Otto 2007; Meirmans & Kolar 2025).

One influential approach frames genome size from a material perspective. Because DNA is phosphorus-rich, variation in nuclear DNA content has been proposed to impose measurable phosphorus costs at cellular and organismal levels (Hessen et al. 2010). Under phosphorus limitation, selection may favor reduced genome size if P allocated to DNA competes with P required for RNA and growth machinery (Hessen et al. 2010). Empirical work in both animals and plants shows that ploidy can influence performance under P limitation and, in some cases, bulk tissue %P (Neiman et al. 2013; Guignard et al. 2016). However, broader comparative analyses indicate that relationships between genome size and tissue N or P concentrations are often weak or context dependent (Bitomsky et al. 2022; Morton et al. 2024), suggesting that genome size effects may be mediated through allocation tradeoffs and physiological fluxes rather than simple proportional increases in bulk P content.

The ionome, defined as the relative abundance of multiple chemical elements within a biological system, provides a natural phenotype for capturing such multielemental organization (Lahner et al. 2003; Salt et al. 2008). Because elemental composition is compositional, changes in one element necessarily imply compensatory shifts in others. Ecological stoichiometry formalizes this mass-balance logic and emphasizes that elemental allocation reflects the combined effects of physiological regulation and environmental supply (Sterner & Elser 2002). Analyses that treat elements independently risk obscuring coordinated allocation patterns that emerge from these constraints. Genome size variation should therefore be expected to reorganize elemental allocation across the organism rather than simply increase demand for any single element.

Compositional data analysis provides an appropriate statistical framework for such systems by accounting for closure and enabling inference on relative allocation among elements (Aitchison 1986; Greenacre 2021). When total organismal chemistry is incompletely observed, the sum of measured elements can be treated as a filling value defining the observed compositional system and making explicit the unmeasured remainder (see Box 1). This approach permits robust comparison of multielemental allocation even when absolute mass or total chemistry are partially unknown, a common limitation in organismal ionomic datasets.

What, then, should genome size variation affect within a multielemental biological system? A common expectation is that effects should be most evident in metabolically active tissues, where changes in transcriptional demand, cell size, and biosynthetic scaling are expressed directly (Bennett 1972; Gregory 2005; Otto 2007; Itgen et al. 2022). These processes rely heavily on coordinated elemental fluxes, particularly of phosphorus-rich nucleic acids and metal-dependent enzymes (Sterner & Elser 2002; Williams & Rickaby 2012), suggesting that genome size effects might be detectable as shifts in elemental allocation within soft tissues.

However, metabolically active tissues are also subject to strong homeostatic regulation. Transport, storage, and turnover mechanisms can buffer elemental composition against environmental variability and underlying genomic differences, potentially dampening cross-ploidy contrasts at steady state (Salt et al. 2008; Jeyasingh et al. 2014; Baxter 2015). An alternative possibility is that genome size effects are expressed more clearly in metabolically inert or slowly turning-over tissues. Mineralized structures such as shells are formed through regulated ion transport and deposition but, once deposited, are largely decoupled from short-term metabolic regulation (Weiner & Dove 2003). Even subtle, persistent differences in elemental allocation may therefore accumulate in such structures over development.

These contrasting expectations motivate a simple question: are genome size effects on elemental composition expressed primarily through short-term metabolic regulation or through long-term integration in mineralized tissue? Addressing this question requires a system in which genome size variation is discrete, tissues differ sharply in metabolic activity, and multielemental composition can be quantified.

The New Zealand freshwater snail *Potamopyrgus antipodarum* provides such a system. Natural populations harbor diploid, triploid, and tetraploid lineages (Neiman et al. 2011) and exhibit pronounced compartmentalization between metabolically active body tissue and a mineralized shell. Here, we quantify ionomes of both tissues and analyze composition using log-ratio methods that explicitly accommodate compositional constraints (Aitchison 1986; Greenacre 2021). By focusing on coordinated shifts in multielemental allocation rather than isolated elemental changes, we test whether ploidy reshapes elemental composition more strongly in metabolically active tissue or in mineralized structure, thereby clarifying how genome size and ionomes are associated.

## Materials and Methods

### Study system and sampling design

Lab-reared isofemale lineages of snails were established from field-collected individuals sampled from two South Island, New Zealand lakes. These lakes, Mapourika and Selfe, were expected to produce a fairly even mix of both diploid and triploid individuals (Neiman et al. 2011, Paczesniak et al. 2013). We established three replicates from different lineages for each combination of lake of origin, ploidy, and ontogenetic stage. Lineages were housed individually under standard laboratory conditions and fed dried *Spirulina* and calcium supplements in excess.

### Sampling ontogenetic stage

We sampled the first ontogenetic stage immediately after juvenile snails emerged from their ovoviviparous mothers, before the juvenile’s shells began to calcify. At this stage, snails are too small to accurately sex, but we used only female snails for the second and third ontogenetic stage since asexual *P. antipodarum* lineages consist almost entirely of females (Neiman et al. 2012) and ionomes can differ drastically between males and females of the same species (Goos et al. 2017). When the remaining snails in the lineage reached 2 mm in length from shell tip to aperture, we sampled the second ontogenetic stage. Upon reproduction signaling sexual maturity, we sampled the third ontogenetic stage. Sexual females were housed with males from their lineage after the second ontogenetic stage to allow for fertilization. To reach the tissue volume necessary in each sample for ion detection levels, we pooled ten individuals for the first ontogenetic stage, five individuals for the second ontogenetic stage, and used a single individual for the third ontogenetic stage.

### Ploidy determination

F1 offspring of each isofemale lineage were sexed under the microscope. If the male:female ratios were roughly 50/50, corresponding with predicted sex ratios for diploid sexuals, the lineage was considered as such (Krist et al., 2014). If ploidy could not be determined by sex ratios, we used flow cytometry to compare sample DNA content with standards of known ploidy. For each lineage in question, an individual was dissected, snap-frozen in liquid nitrogen and prepared for flow cytometry following Krist et al. (2014). Flow cytometry was performed on the Becton Dickinson LSR at the Carver College of Medicine’s Flow Cytometry facility. Ploidy was determined by analyzing peak fluorescence values of stained DNA using FloJo software (following, e.g., Neiman et al. 2011).

### Specimen Preparation

We fasted adult female snails to reduce sample contamination from undigested food. We recorded the wet mass of each individual, then dissected snails into body and shell components. We separated body tissue and shell fragment into individual microcentrifuge tubes, which were snap-frozen in liquid nitrogen, then stored at -80°C. We ultimately used 124 snails from lake Mapourika (62 body, 62 shell samples) and 84 snails from Selfe n = 84 (42 body, 42 shell). Snails belonged to two ploidy levels: 2n n = 132 (66 body, 66 shell); 3n n = 76 (38 body, 38 shell). For each individual, metabolically active soft tissue (body) and mineralized shell were processed separately (n = 208 tissue samples total; n = 104 body, n = 104 shell). Ploidy (2n vs 3n) and tissue type (body vs shell) were treated as focal biological factors impacting the ionome.

### Sample processing and elemental analysis

Snail body and shell tissues were processed separately for elemental analysis. Soft tissues and shells were dissected, dried to constant mass, and digested in trace-metal–grade nitric acid (HNO_3_) following established ionomic protocols. Digestions were conducted for a minimum of 72 h to ensure complete dissolution of organic material and mineral phases, after which samples were diluted to a known final volume using ultrapure (Type I) water.

Elemental concentrations were quantified using inductively coupled plasma–optical emission spectroscopy (ICP-OES; iCAP 7400, Thermo Scientific, Waltham, MA). A broad multielement panel was measured, including major ions and trace elements relevant to physiological regulation and biomineralization (Al, As, B, Ba, Ca, Cd, Co, Cr, Cu, Fe, K, Li, Mg, Mn, Mo, Na, Ni, P, Pb, S, Se, Si, Sr, V, Zn). Multielement external calibration standards (CPI International, Santa Rosa, CA) were used to calibrate the instrument across the expected concentration ranges, and an inline Yttrium internal standard (Peak Performance Inorganic Y Standard, CPI International) was used to correct for instrument drift and matrix effects. Procedural blanks were prepared by digesting acid without tissue material and processed alongside samples to account for background concentrations. Elements that were inconsistently detected across samples were retained in the raw dataset but excluded from multivariate analyses where appropriate, as described below.

Elemental concentrations are reported in solution units. Sample mass was unavailable or heterogeneous for a subset of samples, precluding consistent normalization to dry mass. Accordingly, subsequent analyses were designed to quantify relative elemental allocation using log-ratio transformations anchored to the measured elemental pool, rather than absolute elemental content (Greenacre 2021).

### Filling value and denominator definition

For each sample, we defined a filling value (Fv) as the sum of measured elemental concentrations across a core set of elements that were consistently detected across samples. The core panel comprised 13 elements with ≤5% missing values (B, Ca, Cu, Fe, K, Li, Mg, Mn, Na, P, S, Si, Zn). This core filling value represents the size of the reliably observed elemental pool for each sample. Importantly, the filling value does not assume that all organismal chemistry is measured. Instead, it explicitly conditions inference on the measured chemical pool and treats unmeasured chemistry as an unknown remainder. Negative elemental concentrations were replaced with element-specific limits of detection (LOD), defined as the minimum positive value observed for each element, prior to log-ratio transformation.

### Statistical framework

To test whether metabolically active soft tissue and mineralized shell occupy distinct multielemental regimes, we analyzed elemental composition using log-ratio methods that respect compositional constraints. Elemental data are inherently compositional because each observation represents relative parts of a constrained whole. Analyses based on raw concentrations can therefore generate spurious correlations when denominator variation is ignored.

To test whether genome size variation reorganizes elemental allocation, we computed additive log-ratios (ALRs) for all measured elements, using the core filling value as the explicit denominator. For each element *i*, the ALR was defined as:

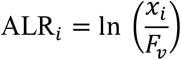

where *x*_*i*_ is the measured concentration of element *i* and *F*_*v*_is the core filling value for that sample. This transformation expresses the relative allocation of each element conditional on the total measured elemental pool and is invariant to unknown multiplicative scaling factors such as sample mass or dilution.

To test whether ploidy effects differ between metabolically active and inert tissues, we fit multivariate models including tissue type, ploidy, their interaction, lake, and developmental stage. We then conducted tissue-specific analyses to evaluate ploidy effects within soft tissue and shell separately while controlling for environmental and ontogenetic context.

Statistical analyses were conducted using multivariate and univariate models applied to ALR-transformed data. Multivariate differences in ionomic composition were evaluated using MANOVA, with Wilks’ lambda used for inference. We diagnosed ploidy-associated elemental imbalances using ALRs anchored to the measured elemental pool, analogous to imbalance analyses previously used to quantify deviation from ancestral ionomic states (Jeyasingh et al. 2023). Diploid (2n) *P. antipodarum* represent the ancestral form (e.g., Neiman et al. 2011) and were thus treated as the reference state, such that ploidy effects represent log-ratio deviations in elemental allocation in triploids relative to diploids.

## Results

### Ionomic quantification

Of the 25 elements measured, 13 (B, Ca, Cu, Fe, K, Li, Mg, Mn, Na, P, S, Si, Zn) exhibited ≤5% missing values and were retained as a core ionomic panel for multivariate analyses. Twelve elements (Al, As, Ba, Cd, Co, Cr, Mo, Ni, Pb, Se, Sr, V) with high (>25%) missingness (i.e., below instrument LOD or true absence) were excluded from multivariate inference to avoid uneven sample representation and distortion of compositional covariance structure. When analyzed together, ionomic data showed hierarchical structuring, with ploidy remaining a consistent effect (Box 2).

### Tissue defines distinct regimes of elemental allocation

Elemental allocation differed strongly between metabolically active soft tissue and mineralized shell (Figure 1). In a reduced multivariate model including tissue type, ploidy, and their interaction, tissue type explained a large proportion of multielemental variation in ALR space (Table 1). This effect was independent of genome size and indicates that soft tissue and shell occupy distinct ionomic regimes defined by relative elemental allocation rather than absolute concentrations.

**Table 1.**
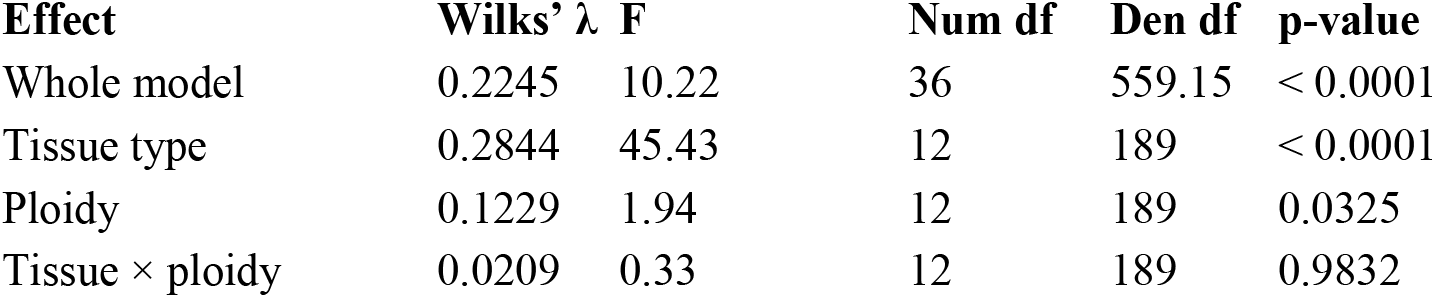
Multivariate test statistics for tissue and ploidy. Multivariate tests were conducted on additive log-ratios (ALRs; ln(element | Fv)) using MANOVA. Wilks’ lambda (λ), approximate F-statistics, numerator and denominator degrees of freedom (df), sample size (n), and associated p-values are reported. Analyses were conducted on 13 ALRs corresponding to elements with ≤5% missing values. The model includes tissue type, ploidy, and their interaction.

**Fig. 1.**
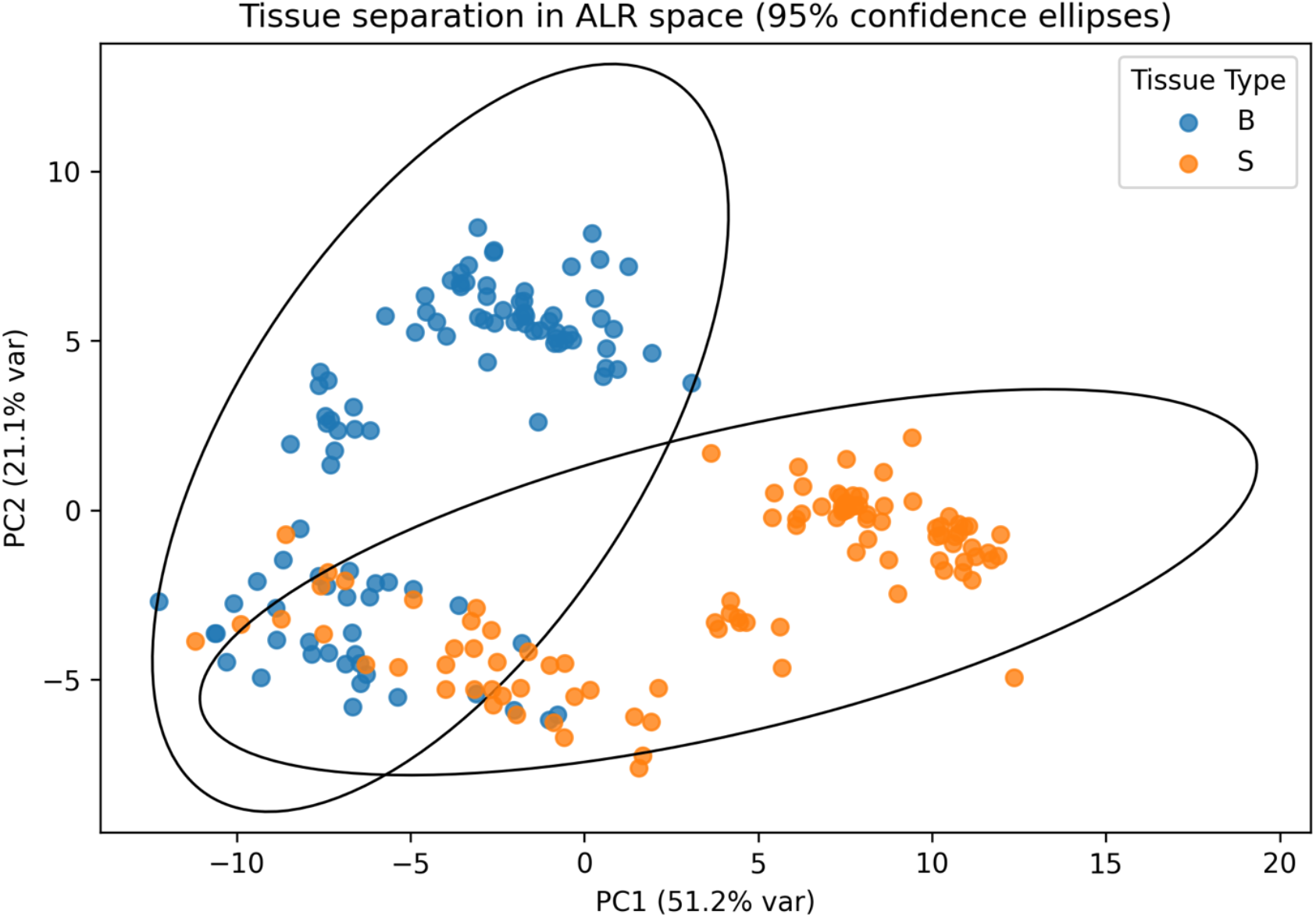
Tissue defines distinct regimes of elemental allocation. Additive log-ratios (ALRs; ln(element | Fv)) reveal strong differences in relative elemental allocation between metabolically active soft tissue and mineralized shell in *Potamopyrgus antipodarum*. Values represent means ± SE across all samples. In ALR space, shells show higher relative allocation to mineral-associated elements (e.g. Ca), whereas soft tissues show higher relative allocation to elements associated with ionic and metabolic regulation (e.g. Na, K, Mg, P). Tissue type explained a large proportion of multielemental variation in ALRs (MANOVA, Wilks’ λ = 0.431, F = 20.87, *p* < 0.0001).

### Ploidy reorganizes relative elemental allocation

Across tissues, ploidy was associated with significant multielemental differences in ALRs, indicating that genome size variation reorganizes relative elemental allocation within the measured elemental pool (Table 1). Although weaker than the tissue effect, this result demonstrates a detectable, system-wide influence of genome size on elemental allocation conditional on Fv. The interaction between tissue type and ploidy was not supported (Table 1), indicating that ploidy-associated ionomic reorganization is broadly consistent across metabolically active and mineralized tissues at the multivariate level.

Ploidy-associated elemental imbalances were further resolved at the level of individual elements using tissue-specific linear models in ALR space (Figure 2). When expressed as ln(element | Fv), ploidy effects were centered on the diploid (2n) reference state, allowing direct interpretation of positive and negative departures in relative elemental allocation. In both tissues, ploidy effects were heterogeneous across elements, indicating selective reorganization of elemental fractions rather than uniform scaling of the ionome.

**Fig. 2.**
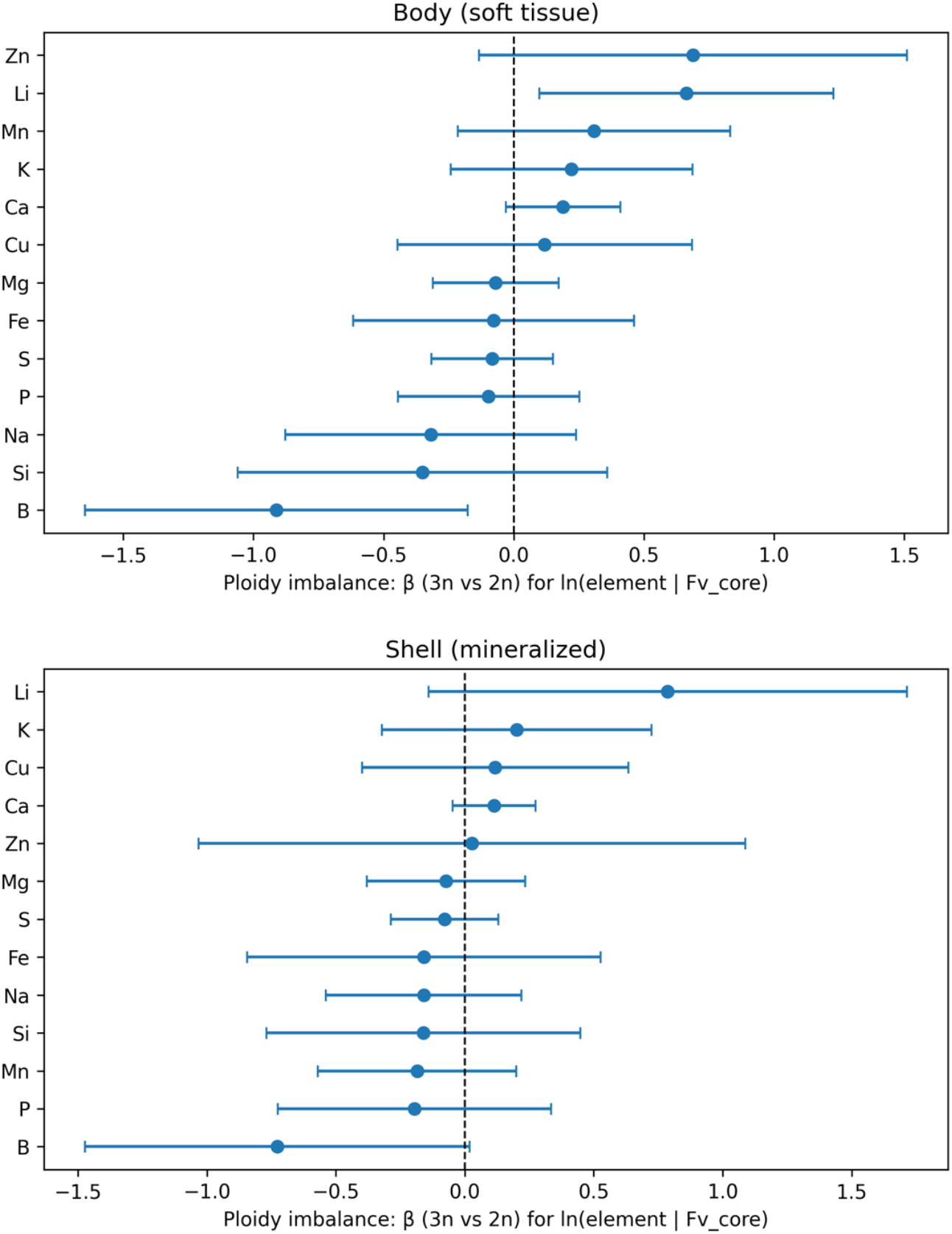
Tissue-specific elemental imbalances associated with ploidy. Ploidy-associated elemental imbalances were quantified using additive log-ratios anchored to the measured elemental pool (ln(element | Fv)), where Fv is the sum of reliably measured elements. Panels show results for (A) metabolically active soft tissue and (B) mineralized shell. Points indicate estimated ploidy effects (β, 3n vs 2n) from linear models fit separately for each element within tissue, controlling for lake and developmental stage; error bars show 95% confidence intervals. The vertical dashed line at zero represents the diploid (2n) reference state. Values less than zero indicate elements that comprise a smaller fraction of the measured elemental pool in triploids (3n) relative to diploids, whereas values greater than zero indicate elements that comprise a larger fraction of the measured elemental pool in triploids. Confidence intervals that do not overlap zero indicate statistically detectable ploidy-associated imbalance in relative elemental allocation.

In metabolically active soft tissue, ploidy-associated imbalances were generally modest, with fewer elements exhibiting confidence intervals that excluded zero (Figure 2A). By contrast, mineralized shell showed stronger and more frequent departures from the diploid reference state (Figure 2B), consistent with the larger multivariate ploidy effect detected for shell ALRs. Elements with confidence intervals not overlapping zero indicate statistically detectable ploidy-associated shifts in their proportional contribution to the measured elemental pool, conditional on Fv and after accounting for lake and developmental stage.

Together, these tissue-specific imbalance profiles demonstrate that genome size effects on elemental allocation are expressed asymmetrically across biological compartments, with mineralized structures showing greater sensitivity to ploidy than metabolically active soft tissue.

## Discussion

Genome size variation is often invoked as a determinant of organismal performance, yet the material pathways linking genome architecture to phenotype remain difficult to resolve, particularly in animals. By analyzing elemental composition as a compositional, multielemental phenotype, we show that ploidy reshapes organismal chemistry through coordinated redistribution of elements rather than through changes in isolated elemental demands. These effects are tissue structured and element specific, revealing how genome size variation propagates through biological compartments with distinct regulatory and integration timescales.

### Tissue structure defines the chemical backdrop for ploidy effects

Tissue type overwhelmingly structured ionomic composition (Fig. 1), confirming that metabolically active soft tissue and mineralized shell occupy distinct chemical regimes. This partition reflects fundamental differences in turnover rate, regulatory control, and developmental integration between tissues (Sterner & Elser 2002; Weiner & Dove 2003). Genome size effects therefore act on a chemically compartmentalized organism rather than on a homogeneous pool.

### Genome size reorganizes elemental allocation, not absolute elemental quotas

Although weaker than tissue effects, ploidy exerted a statistically detectable multivariate influence on elemental allocation (Table 1). Importantly, this effect was expressed as redistribution across multiple elements rather than as strong shifts in any single element. This pattern is inconsistent with simple phosphorus-cost hypotheses that predict proportional increases in P demand with increased genome size due to the P-rich nature of nucleic acids (Hessen et al. 2010). Instead, genome duplication appears to reorganize the relative partitioning of elements within the measured pool. Similar element-specific rather than phosphorus-dominated shifts have been reported in diploid versus autotetraploid plants, where genome duplication altered Fe, Ca, and Zn allocation without consistent increases in bulk %P (Simko et al. 2023).

Previous work in *Potamopyrgus* showed that ploidy alters performance under phosphorus limitation and can influence P use and excretion (Neiman et al. 2013). The present analysis addresses a complementary question: whether ploidy reorganizes steady-state elemental allocation across a multielement system. The absence of strong bulk %P differences here does not contradict earlier findings; rather, it indicates that genome size effects may be expressed through coordinated redistribution or altered fluxes that are not detectable through single-element concentration alone.

A recent whole-genome assembly for *P. antipodarum* documents a relatively recent whole-genome duplication with substantial duplicate retention (Jalinsky et al. 2025). Expansion of gene copy number increases the potential landscape of structural macromolecules, transporters, and metal-binding proteins, even when expression is buffered. Such expansion provides a mechanistic basis for subtle shifts in uptake, binding, compartmentalization, and deposition probabilities across elements. Under this view, ploidy-associated ionomic differences reflect altered system-level allocation dynamics rather than fixed increases in demand for any specific element.

### Tissue-specific expression of ploidy effects

Ploidy-associated imbalances were modest in metabolically active soft tissue but more pronounced in mineralized shell (Fig. 2). Soft tissues are subject to active regulation of uptake, storage, and turnover, which can buffer steady-state composition despite underlying genomic differences (Salt et al. 2008; Jeyasingh et al. 2014). In contrast, shells integrate elemental deposition over extended developmental timescales and are largely decoupled from short-term metabolic regulation (Weiner & Dove 2003). Even small, persistent differences in transport or incorporation rates can therefore accumulate into detectable compositional shifts.

Elements with no established biochemical role in animals, notably lithium and boron, showed some of the clearest ploidy-associated signals. Because these ions interact primarily through weak substitution and non-specific binding (Williams & Fraústo da Silva 1996, 2002), their redistribution is unlikely to reflect targeted physiological demand. Instead, they may act as sensitive reporters of changes in the physical and chemical architecture of cells and tissues associated with genome duplication. This interpretation reinforces the view that ploidy reshapes elemental organization indirectly, through changes in binding landscapes and transport probabilities rather than through selection on simple stoichiometric requirements.

### Implications

Our analysis prioritizes tractability over chemical completeness. We did not quantify bulk constituents such as C, H, O, and N, nor elements that are difficult to measure reliably with ICP-based methods (e.g. chloride or ultra-trace metals). The filling value therefore represents the measured elemental pool rather than total organismal chemistry. By expressing composition as log-ratios anchored to this pool, inference is explicitly conditioned on observed chemistry while unmeasured components remain an unspecified remainder (Aitchison 1986; Greenacre 2021).

Taken together, these results show that genome size variation leaves a coherent multielemental signature in organismal chemistry, even when single-element effects are weak. Ionomic composition therefore provides a sensitive phenotype for detecting the material consequences of genome duplication. By focusing on relative allocation within a measured elemental pool, compositional analysis reveals how genome architecture propagates through rates of uptake, retention, and deposition to shape tissue-level chemical organization. More broadly, this approach offers a new way to investigate ploidy and genome size: not only through gene content or performance traits, but through system-level patterns of elemental allocation that integrate genomic, physiological, and environmental influences.

## Acknowledgements

We thank anonymous reviewers for constructive feedback. This work was partially supported by Iowa Office for Undergraduate Research and the Linda and Rick Maxson Undergraduate Research Award. We would also like to acknowledge The University of Iowa Biology Honors Program and Lori Adams for providing support to MR. Elemental analyses were supported by institutional overhead return funds allocated to PDJ by the College of Arts & Sciences at Oklahoma State University from prior externally funded awards. YI was supported by a Grand River Dam Authority Graduate Research Fellowship. Generative AI was used to assist with language editing for clarity in preparation of this manuscript.

## Figures, Tables, Boxes

### Box 1. Genome size, ionomes, and organisms as dynamical material systems far from equilibrium

Organisms are open material systems maintained far from equilibrium by continuous fluxes of chemical elements. At any moment, the elemental composition of a tissue reflects the balance of elemental uptake, transport, storage, transformation, and loss. Genome size variation, such as changes in ploidy, is therefore expected to influence elemental composition indirectly by altering the rates of these underlying processes rather than by changing elemental requirements in isolation.

Let x_i_(t) denote the amount of element *i* in a given tissue at time *t*. The dynamics of elemental composition can be represented schematically as:

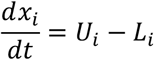

where U_i_ represents the aggregate rate of uptake and internal allocation of element *i*, and L_i_ represents the aggregate rate of loss via excretion, turnover, or sequestration into other compartments. Genome size can modify these rates by altering cell size, transcriptional demand, surface-to-volume ratios, and the scaling of transport and biosynthetic machinery, even when external elemental supply remains constant.

Because elemental data are compositional, biological inference depends on relative allocation rather than absolute quantities. We therefore express elemental composition using additive log-ratios (ALRs) relative to the filling value (Fv), defined as the sum of reliably measured elements:

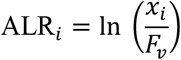

This transformation reframes elemental composition as a state variable describing how each element is partitioned within the measured elemental pool, independent of unknown multiplicative factors such as tissue mass or dilution. Changes in ALRs therefore indicate redistribution of elemental fractions driven by differences in underlying rates, not simple concentration effects.

Tissue type determines how these rate processes are expressed. Metabolically active soft tissues are characterized by rapid turnover and strong homeostatic regulation, which can buffer short-term perturbations in elemental fluxes. In contrast, mineralized structures such as shells integrate elemental allocation over longer timescales, effectively accumulating the net outcome of small, persistent biases in uptake, transport, or loss. As a result, genome size effects that weakly influence instantaneous metabolic rates may nonetheless produce pronounced compositional signatures in inert tissues.

From this perspective, ploidy does not impose a fixed elemental “signature” but instead reshapes the dynamical landscape of elemental fluxes. Ionomic differences between diploid and triploid organisms thus reflect altered trajectories through compositional space, revealing how genome architecture propagates upward to cellular and organismal material composition through changes in rates rather than static requirements.

### Box 2. Hierarchy of biological variation in *Potamopyrgus antipodarum* ionomes

Ionomic composition reflects multiple nested sources of variation operating at distinct biological scales. Multivariate analysis of 13 additive log-ratios (ALRs; N = 204) reveals a clear hierarchy of effects.

#### 1. Developmental stage

Developmental stage exerts the strongest multivariate influence on ionomic composition (MANOVA, F_12_,_188_ = 92.89, p < 0.0001). This large effect reflects ontogenetic changes in traits and associated elemental signatures (e.g., Back & King 2013; Sherman & Jeyasingh, in revision; Loganathan & Jeyasingh, in review), and is consistent with other data demonstrating distinctly different body composition in juvenile and adult *P. antiopodarum* (Najev et al. 2026, in press). Stage was included to prevent confounding ploidy with ontogenetic scaling.

#### 2. Tissue compartment

Tissue type (soft tissue vs shell) is also a dominant determinant of elemental allocation (F_12_,_188_ = 45.45, p < 0.0001). This strong partition reflects fundamental differences in turnover rate, integration timescale, and biomineralization processes. Tissue establishes the primary structural background against which other effects are expressed.

#### 3. Lake of origin

Lake contributes a significant but smaller multivariate effect (F_12_,_188_ = 7.17, p < 0.0001), consistent with differences in environmental elemental supply previously demonstrated for the New Zealand lakes that harbor *P. antipodarum* (Krist et al. 2016). Because only two lakes were sampled, lake is treated as background geochemical context rather than as a focal ecological variable.

#### 4. Genome size

Conditional on tissue, developmental stage, and lake, ploidy retains a statistically significant multivariate effect (F_12_,_188_ = 2.30, p = 0.0093). Although smaller in magnitude than other sources in explaining ionomic variance, this effect emerges as coordinated redistribution across elements in ALR space and is rarely detectable at the single-element level. The persistence of this signal after conditioning on strong ontogenetic and structural partitioning indicates that genome size subtly but coherently reshapes relative elemental allocation.

#### Boundary of inference

This study isolates genome size (ploidy) as the primary axis of interest while statistically conditioning on tissue structure, developmental scaling, and geochemical background. Developmental stage and tissue identity explain the majority of total multielemental variance, as expected for growth and compartmental partitioning. By contrast, the ploidy effect is smaller but of general significance: it represents a persistent redistribution within compositional space that cannot be attributed to ontogeny or environmental context. Because whole-genome duplication is a widespread and recurrent feature of eukaryotic evolution, understanding how genome size reshapes multielement organization has general relevance beyond this system. The present analysis therefore frames genome size as a broadly applicable material impulse expressed against a hierarchically structured organism.

## Files for Electronic Supplement

**Supplementary Data S1**. Mass-specific elemental concentrations and metadata for *Potamopyrgus antipodarum* ionome analyses. Merged dataset containing sample metadata (lake, ploidy, developmental stage, lineage, tissue type) and elemental concentrations for all measured elements. Concentrations are provided in solution units (µg/L). The full elemental panel (sheet: Raw) is included prior to selection of the core subset (sheet: Curated) used in transformations and subsequent multivariate analyses.

## References

Aitchison J. 1986 The statistical analysis of compositional data. London, UK: Chapman & Hall.

Back, J. A., and R. S. King. 2013. Sex and size matter: ontogenetic patterns of nutrient content of aquatic insects. Freshwater Science 32:837–848.

Baxter I. 2015 Should we treat the ionome as a combination of individual elements, or should we be deriving novel combined traits? J Exp Bot 66, 2127–2131. (doi:10.1093/jxb/erv040)

Bennett MD. 1972 Nuclear DNA content and minimum generation time in herbaceous plants. Proc R Soc Lond B 181, 109–135.

Bitomsky M, Kobrlová L, Hroneš M, Klimešová J, Duchoslav M. 2022 Stoichiometry versus ecology: relationships between genome size, GC content, and tissue nitrogen and phosphorus in grassland herbs. Ann Bot 130, 189–197. (doi:10.1093/aob/mcac079)

Cavalier-Smith T. 2005 Economy, speed and size matter: evolutionary forces driving nuclear genome miniaturization and expansion. Ann Bot 95, 147–175.

Elser JJ, Hamilton A. 2007 Stoichiometry and the new biology: the future is now. PLoS Biol 5, e181. (doi:10.1371/journal.pbio.0050181)

Gregory TR. 2005 Genome size evolution in animals. In The evolution of the genome (ed TR Gregory), pp. 3–87. San Diego, CA: Elsevier.

Greenacre M. 2021 Compositional data analysis. Annu Rev Stat Appl 8, 271–299.

Guignard MS et al. 2016 Impacts of whole-genome duplication on plant growth and elemental stoichiometry. New Phytol 210, 262–273.

Hessen, D. O., P. D. Jeyasingh, M. Neiman, and L. J. Weider. 2010. Genome streamlining and the elemental costs of growth. Trends in ecology & evolution 25:75–80.

Itgen MW, Natalie GR, Siegel DS, Sessions SK, Mueller RL. 2022 Genome size diversity in salamanders is driven by phylogenetic constraints and ecological variables. Genome Biol Evol 14, evac105.

Jalinsky, J., K. E. McElroy, J. Sharbrough, L. Bankers, P. D., C. Higgins, C. Toll, et al. 2025. Whole-genome sequence of Potamopyrgus antipodarum-A model system for the maintenance of sexual reproduction-reveals a recent whole-genome duplication. Genome Biology and Evolution 17:evaf192.

Jeyasingh PD, RD Cothran, M Tobler (2014). Testing the ecological consequences of evolutionary change using elements. Ecology & Evolution 4: 528–538.

Jeyasingh PD, P Roy Chowdhury, MW Wojewodzic, D Frisch, DO Hessen, LJ Weider (2015). Phosphorus use, and excretion varies with ploidy level in Daphnia. Journal of Plankton Research 37: 1210–1217.

Jeyasingh PD, R Sherman, C Prater, K Pulkkinen, T Ketola (2023). Adaptation to a limiting element involves mitigation of multiple elemental imbalances. Journal of the Royal Society Interface 19: 20220472.

Krist AC, Kay AD, Larkin K, Neiman M. 2014 Response to phosphorus limitation varies among lake populations of the freshwater snail Potamopyrgus antipodarum. PLoS ONE 9, e85845.

Krist AC, Kay AD, Scherber E, Larkin K, Brown BJ, Lu D, Warren DT, Riedl R, Neiman M. 2016 Evidence for extensive but variable nutrient limitation in New Zealand lakes. Evol Ecol 30, 973–990.

Lahner B et al. 2003 Genomic scale profiling of nutrient and trace elements in Arabidopsis thaliana. Nat Biotechnol 21, 1215–1221.

Leitch IJ, Leitch AR. 2008 Genomic plasticity and the diversity of polyploid plants. Science 320, 481–483.

Lotka AJ. 1925 Elements of physical biology. Baltimore, MD: Williams & Wilkins.

Meirmans, P. G., and F. Kolar. 2025. Whole-genome duplication leads to significant but inconsistent changes in climatic niche. Proceedings of the National Academy of Sciences of the United States of America 122:e2424785122.

Morton, J. A., C. A. Arnillas, L. Biedermann, E. T. Borer, L. A. Brudvig, Y. M. Buckley, M. W. Cadotte, et al. 2024. Genome size influences plant growth and biodiversity responses to nutrient fertilization in diverse grassland communities. PLoS Biology 22:e3002927.

Najev BL, Gavin WG, Craven C, Escandón A, Pate P, Krist AC, Neiman M. 2026 Phosphorus availability affects multiple metrics of reproductive investment in a freshwater snail. Royal Society Open Science, in press

Neiman M, Paczesniak D, Soper DM, Baldwin AT, Hehman G. 2011 Wide variation in ploidy level and genome size in a New Zealand freshwater snail with coexisting sexual and asexual lineages. Evolution 65, 3202–3216.

Neiman M, Larkin K, Thompson AR, Wilton P. 2012 Male offspring production by asexual Potamopyrgus antipodarum, a New Zealand snail. Heredity 109, 57–62.

Neiman M, Kay AD, Krist AC. 2013 Sensitivity to phosphorus limitation increases with ploidy level in a New Zealand snail. Evolution 67, 1511–1517.

Otto SP. 2007 The evolutionary consequences of polyploidy. Cell 131, 452–462.

Paczesniak D, Jokela J, Larkin K, Neiman M. 2013 Discordance between nuclear and mitochondrial genomes in sexual and asexual lineages of the freshwater snail Potamopyrgus antipodarum. Mol Ecol 22, 4695–4710.

Roddy AB et al. 2020 Scaling of genome size and cell size limits maximum photosynthetic rates. New Phytol 225, 1216–1229.

Salt DE, Baxter I, Lahner B. 2008 Ionomics and the study of the plant ionome. Annu Rev Plant Biol 59, 709–733.

Simko, I., and R. Zhao. 2023. Phenotypic characterization, plant growth and development, genome methylation, and mineral elements composition of neotetraploid lettuce (Lactuca sativa L.). Frontiers in Plant Science 14:1296660.

Sterner RW, Elser JJ. 2002 Ecological stoichiometry: the biology of elements from molecules to the biosphere. Princeton, NJ: Princeton University Press.

Weiner S, Dove PM. 2003 An overview of biomineralization processes and the problem of the vital effect. Rev Mineral Geochem 54, 1–29.

Williams RJP, Fraústo da Silva JJR. 1996 The natural selection of the chemical elements. Oxford, UK: Oxford University Press.

Williams RJP, Fraústo da Silva JJR. 2002 The biological chemistry of the elements: the inorganic chemistry of life, 2nd edn. Oxford, UK: Oxford University Press.

Williams RJP, Rickaby REM. 2012 Evolution’s destiny: co-evolving chemistry of the environment and life. Cambridge, UK: Royal Society of Chemistry.

